# Strategic adjustment of parental care: life-history trade-offs and the role of glucocorticoids

**DOI:** 10.1101/063313

**Authors:** Çağlar Akçay, Ádám Z. Lendvai, Mark Stanback, Mark Hausmann, Ignacio T. Moore, Fran Bonier

**Affiliations:** Department of Biological Sciences, Virginia Tech, Blacksburg, VA, USA; Department of Evolutionary Zoology and Human Biology, University of Debrecen, Debrecen, Hungary; Department of Biology, Davidson College, Davidson, NC, USA; Department of Biology, Bucknell University, Lewisburg, PA, USA; Department of Biology, Queen’s University, Kingston, ON, Canada

**Keywords:** tree swallow, corticosterone, nestling begging, provisioning, brood value, latitude, fitness

## Abstract

Life history theory predicts that optimal strategies of parental investment will depend on ecological and social factors such as current brood value and offspring need. Parental care strategies are also likely to be mediated in part by the hypothalamic-pituitary-adrenal (HPA) axis and glucocorticoid hormones. Here we present an experiment in tree swallows (*Tachycineta bicolor*), a biparental songbird with wide geographic distribution, asking whether parental care is strategically adjusted in response to signals of offspring need and brood value and whether glucocorticoids are involved in these adjustments. Using an automated playback system, we carried out playbacks of nestling begging calls specifically to females in two populations differing in their brood value: a northern population in Ontario, Canada (relatively high brood value) and a southern population in North Carolina, USA (lower brood value). We quantified female offspring provisioning rates before and during playbacks and plasma corticosterone levels (cort) once during late incubation and once immediately after playbacks. Females in both populations increased feeding rates temporarily during the first two hours of playback but the increase was not sustained for the entire duration of playback (six hours). Cort levels from samples at the end of the playback did not differ between control females and females that received playbacks. However, females that had higher increases in cort between the incubation and nestling period had greater fledging success. These results suggest that females are able to strategically respond to offspring need, although the role of glucocorticoids in this strategic adjustment remains unclear.

## Introduction

Parental investment comprises costly behaviors that can improve the survival of current offspring at the expense of future reproduction (Trivers, 1972). Life history theory predicts that animals will adopt strategies that optimize the level of parental investment in a given reproductive bout. To determine the optimal level of investment, parents must incorporate several potential cues about the value of both current and future reproductive activities (Stearns, 1992). For instance, parents may adjust their investment in offspring in response to cues that indicate genetic quality or condition of their partner (Burley, 1988; Dakin et al., 2016; Sheldon, 2000) or parental effort of their partner (Hinde, 2006; Hinde and Kilner, 2007; Westneat et al., 2011). In addition, parents may also adjust investment based on cues from their offspring such as the frequency and intensity of begging calls and begging displays (Leonard and Horn, 2001; Mock et al., 2011; Ottosson et al., 1997).

Decisions about allocation of parental resources can also depend on life history strategy. In particular, the likelihood of future reproduction is expected to affect the level of parental investment in current reproduction. For instance, in bird populations breeding at more extreme latitudes, the potential for future reproduction tends to be lower due to lower adult survival. Consequently, parents invest more in current reproduction in higher latitudes compared to birds breeding in lower latitudes (Ardia, 2005; Ghalambor and Martin, 2001; Martin et al., 2000).

Resource allocation decisions may be proximately regulated by the physiological state of an individual (McNamara and Houston, 1996). For example, the hypothalamic-pituitary-adrenal (HPA) axis, and glucocorticoid hormones in particular, can mediate the trade-off between resources allocated to reproduction or self-maintenance (Bonier et al., 2009a; Wingfield and Sapolsky, 2003; Wingfield et al., 1998). An acute increase in glucocorticoids might trigger allocation of energetic resources to selfmaintenance and survival at the expense of allocation to reproductive effort (Wingfield and Sapolsky, 2003; Wingfield et al., 1998). Conversely, experimental studies showed that individuals with high current reproductive effort have a reduced glucocorticoid response to stressors (Lendvai and Chastel, 2008; Lendvai et al., 2007). Finally, glucocorticoid levels often covary negatively with measures of individual condition or habitat quality (Moore et al., 2000). Based on these lines of evidence, levels of glucocorticoids are often expected to be negatively correlated with reproductive investment (Bonier et al., 2009a).

More recently, evidence has started to accumulate that corticosterone (the main avian glucocorticoid, henceforth referred to as “cort”) at baseline levels may support reproductive investment such that cort levels actually increase with some metrics of reproduction (Bonier et al., 2009b; Bonier et al., 2011; Crossin et al., 2012; Love et al., 2004; Moore and Jessop, 2003; Ouyang et al., 2013). This hypothesis, termed the Cort-Adaptation Hypothesis, suggests that increased baseline cort levels can induce increased foraging and offspring provisioning behavior and decreased sensitivity to acute stress, which in turn leads to higher reproductive effort and fitness (Angelier and Chastel, 2009; Angelier et al., 2007; Bonier et al., 2009a). Consistent with this hypothesis, brood value and baseline cort are positively correlated across species (Bokony et al., 2009), implying that baseline cort reflects high investment in current reproduction.

Evidence for the Cort-Adaptation hypothesis also comes from studies of variation in parental investment and cort within species. For instance in a recent study of tree swallows (*Tachycineta bicolor*), females with higher baseline cort during the offspring provisioning stage fledged more young than females with lower cort (Bonier et al., 2009b). Female swallows also increased baseline cort levels when they were caring for an experimentally enlarged brood, as compared to females caring for experimentally reduced broods (Bonier et al., 2011). Similarly in house sparrows (*Passer domesticus*), the number of fledglings a female was able to raise was positively correlated with the change in the females’ baseline cort levels from pre-laying to the nestling feeding period (Ouyang et al., 2011). In another study on macaroni penguins (*Eudyptes chrysolophus*), experimentally increased cort levels within the normal range of baseline caused increased foraging activity of females who subsequently raised heavier chicks than control implanted females (Crossin et al. (2012).

Here, we ask two related questions regarding the strategic adjustment of parental care in response to offspring signals in two box-nesting populations of tree swallows in Ontario, Canada and North Carolina, USA. We manipulated offspring demand perceived by females through the use of an automated playback system (Lendvai et al., 2015b) where nestling begging calls were directed specifically to females for six hours when nestlings were six days of age. In addition, we measured baseline cort levels in the same females once during the incubation period and once immediately after the playback period (9-13 days after the first sample, depending on the hatch date).

Tree swallows in our southern population in North Carolina have higher annual return rates – a robust proxy for annual survival in this highly philopatric species (Winkler et al., 2004) – and a longer breeding season that in some instances even allows birds to raise a second brood (MS & ÇA, unpublished data). In contrast, tree swallows in Ontario have lower annual return rates and a shorter breeding season, with only one brood raised per pair each year. Greater annual survival in the North Carolina population means that, on average, these birds have more opportunities for future reproduction than birds in Ontario. Higher potential for future reproduction will tend to decrease the value of the current brood (Ardia, 2005), such that current brood value will be lower for NC tree swallows compared to Ontario tree swallows. Thus, the use of these two populations allows us to compare responsiveness of parental investment to cues of offspring demand across populations with different brood values (Bokony et al., 2009; Silverin et al., 1997; Sol et al., 2012).

The first question we ask is whether adjustment of parental care effort to offspring begging calls differs between the two populations. This question relates to the Brood Value Hypothesis which predicts that adjustment of parental investment in response to cues from offspring will depend on the value of current reproduction vs. future reproduction (Silverin et al., 1997). Thus, this hypothesis predicts that females should increase parental care in response to experimentally increased offspring demand more (or only) in the northern population with higher brood value compared to the southern population.

The second question relates to the role of cort in mediating strategic adjustments in parental effort. The Cort-Adaptation Hypothesis predicts that increases in parental investment should be positively correlated with increases in cort levels (Bonier et al., 2009a). As such, females that received playbacks should show greater increases cort levels than control females, which did not receive playbacks. We also test the prediction from the Cort-Adaptation hypothesis that higher increases in cort during nestling period will predict higher fledging success.

## Material and Methods

### Study site and species

The tree swallow is a widespread secondary cavity nesting species that breeds across a wide range of latitudes from Alaska and Northern Canada to the southern USA. We studied tree swallows at two field sites where they nest in artificial nest boxes: Queens’s University Biological Station, Ontario, Canada (N44°34’2.02”, W76°19’26.036”, 121 m elevation) and near Davidson College, Davidson, North Carolina (NC), USA (N34°31’ 32.34”, W80°52’40”, 240 m elevation). These two sites differ in the length of the breeding season (May to July in Ontario and March to July in NC). Tree swallows have high breeding site fidelity, and so return to the breeding population is often used as a proxy of annual survival (Winkler et al., 2004). In our NC population, return rates are around 50% for females (51% in 2015), higher than the Ontario population (average 22% between 1975-2012, range 10-45%), as has been found in other studies comparing southern and northern populations of tree swallows (Ardia, 2005). The procedures used in the study followed the guidelines for animal care outlined by Animal Behavior Society and Association for the Study of Animal Behavior, and were approved by approved by the Institutional Animal Care and Use Committee at Virginia Tech (#12-020) and the Canadian Wildlife Service (#10771).

### Nest monitoring

We monitored the nests by visiting each nest box weekly until the parents started nest construction, after which point we visited the nest box every three days until an egg was detected. We checked the nest box every day until no new eggs were laid for two days in a row, which indicated that incubation had started. Female tree swallows typically lay 1 egg per day, and begin incubation on the day of laying of the last egg (Winkler et al., 2011). The date of laying of the last egg was considered day 0 of the incubation period. The incubation period typically lasts 14 days, so we checked each incubating nest daily starting from day 12 of the incubation period until all chicks hatched to determine the date of hatching, which was defined as the day when the first chick hatched. Day of hatching was considered day 0 of the nestling period. Throughout the nestling period, we checked the nest at least every 3 days until day 16, at which point we stopped disturbing the nest until day 22 to determine fledging success.

### Banding parents and nestlings

We captured females using box traps at their nest on day 10 of the incubation period to record body measurements (tarsus, wing chord, weight, skull size), collect a blood sample for cort analysis (within 3 minutes of capture), and mark birds with a numbered metal band and a unique passive integrated transponder (PIT) tag that was integrated into a plastic colored leg band (EM4102 tags from IB Technology, UK). Each female was tagged with a red color band/PIT tag. We captured the males at their nest box on day 2 or 3 of the nestling period. We took the same measurements from the males, except that we did not collect blood samples to minimize handling time and capture stress for males. Males were also tagged with a numbered metal band and a blue PIT tag. We report a detailed analysis of male parental behavior as a function of treatment and female behavior elsewhere. We measured tarsus length and weighed nestlings on the afternoon of day 6 and again on day 12 when each nestling received a numbered metal band.

### Playback experiment

We recorded begging calls from 10 nests on the afternoon of day 6 by pointing a Sennheiser ME66/K6 directional microphone attached to a Marantz PMD 660 Solid State recorder into the nest. To initiate nestling begging, we tapped at the nest entrance, which is a similar sound to what the parents make as they land on the nest box. We used the software Syrinx (John Burt, Seattle, WA; www.syrinxpc.com) to create 30 second stimulus files from the recorded begging calls, with a standard call rate that initially was 14 begs/sec that gradually decreased to a constant 4 begs/sec, simulating a natural pattern of begging in which the nestlings beg vigorously immediately upon arrival of the parent and then tail off gradually. The 10 stimulus files were randomly allocated to the treatment nests. We used a radio-frequency identification (RFID) reader (an upgrade of the model described in Bridge and Bonter, 2011), that was obtained from Cellular Tracking Technology, PA, USA) attached to a microcomputer (Raspberry PI) to carry out the playbacks automatically every time the female (but not the male) perched at the nest box entrance. The playback set-up is described in detail elsewhere (Lendvai et al., 2015b). Briefly, we attached an antenna around the entrance hole of the nest box that was connected to an RFID reader. The RFID reader in turn was connected to a Raspberry PI computer which was running a Python script that played back the begging calls for 30 seconds every time the RFID reader detected the female's PIT tag, with the exception of a refractory period of 2 minutes from the start of each playback (to avoid situations where the playback would be triggered when the female left the nest). The playback apparatus was also installed for control nests, but no sound was played. Treatments were allocated to the nests using a randomized block design, to control for seasonal differences.

The playback system was set up in the morning around 7am on day 6 post hatching and stopped approximately 6 hours later when we captured the females in their nest box and obtained a second blood sample for cort analysis. We had 21 control and 15 playback nests in NC and 19 control and 18 playback nests in Ontario. In NC, three of the nests that were intended to be playback nests never received any playbacks due to the failure of the system, and as such they were included in the analysis as control nests, which caused the uneven sample sizes. One additional nest in NC was excluded from analyses because it only received 2.7 hr of playback due to equipment failure halfway through the experiment, making it intermediate to control and playback conditions.

### Blood sampling and hormone assay

We obtained blood samples (approximately 120 μl) by puncturing the brachial vein within 3 minutes of capturing the females to minimize the influence of the stress of capture on measured cort levels (Romero and Reed, 2005). Blood was stored on ice in the field, and centrifuged in the laboratory within 6 hours to separate the plasma. The plasma was then stored at −20°C until taken to Virginia Tech for hormone assay.

Total cort in plasma was determined by direct radioimmunoassay following an extraction with dichloromethane (Bonier et al., 2009a; Wingfield et al., 1992). Mean extraction efficiency of a known quantity of radiolabeled hormone was 83%, and we corrected for the individual extraction efficiencies in calculating final concentrations. Briefly, we incubated the extracts overnight at 4°C with 10K cpm of 3H-Cort (Perkin Elmer, Product number: NET399250UC) and antiserum (Esoterix Endocrinology, Calabasas Hills, CA 91301, Product number: B3-163). We then added dextran-coated charcoal to separate cort bound to antibodies. Intra-assay variation of known concentration standards was 3.93%.

### Quantifying parental effort

We quantified parental visit rates in two ways: first, we carried out 1-hour feeding watches on day 5 and day 6 (the day before and the day of the treatments) where an observer sat 30 m from the nest and noted every visit of the male and female using a spotting scope and a voice recorder. We also quantified visit rates from the RFID records as described in detail in Lendvai et al. (2015a). We checked the visit rates from 1-hour nest watches against the visit rates calculated from RFID logs of the same time periods. There was a high correspondence between the two (r= 0.68, p=0.2 × 10^−7^, for females and r=0.67, p=0.4 × 10^−7^ for males, n= 43). Because the RFID observations spanned the entire duration of the experiment we used these as the main measure of parental visit rates. Visit rates are an excellent measure of the feeding rates in the tree swallows, as most visits are for feeding (McCarty, 2002).

### Data analyses

We used generalized linear mixed models (GLMM) to assess the effects of treatment and population on cort and behavioral data. We entered the cort data into a GLMM with time period (incubation or nestling), treatment (playback vs. control), and population (Ontario vs. NC) as fixed factors. Playback stimulus and bird ID were included as random factors.

We analyzed the female feeding rates derived from RFID recordings with GLMMs using the fixed factors treatment (playback vs. control), population (Ontario vs. NC), and time period. For the latter factor, we used four levels: pre-treatment (day 5) feeding rates (6 hrs during the same time of day as the experimental period on the next day) and feeding rates from the period while the playback or control treatment was in effect in day 6, which we further divided into three two-hour periods to assess any temporal changes in effects of playback on female behavior. We included playback stimulus and bird ID as random factors and also included an offset variable for log of duration of playback to control for the variation in how long the birds were exposed to the playbacks (mean= 6.24 ± 0.05 SE hours). Because of the large number of predictor variables (3 fixed factors and their interactions) in these mixed models, we used a model averaging approach. We first ran a model with all predictor variables and their interactions and subsequently used model averaging with R-package MuMIn (Bartoń, 2013). In the averaged models, we included all models within 2 AICc of the best model (i.e. the model with the lowest AICc).

We also examined whether change in cort (base-10 logarithm of ratio of post-treatment cort to pre-treatment cort) between incubation and nestling periods predicted number of chicks fledged using a generalized linear model with Poisson distribution and log link (Bonier et al., 2011). For this analysis we included change in cort, treatment, population and their interactions as predictor variables and again used model averaging as above.

Finally, we analyzed nestling growth (nestling mass on day 6 and day 12) as well as fledging success using GLMMs. In these models, we examined the fixed factors population, treatment, and their interactions and included relative lay date (number of days from the first egg of the respective population) as a random factor as it has a strong effect on clutch size with later clutches containing fewer eggs in both populations.

## Results

*Female feeding rates:* Nestling playbacks had a transient effect on female feeding rates. Females that received playbacks of nestling begging calls increased their feeding rates in the first two hours of the playbacks on day 6, compared to their average feeding rates the previous day. No such increase was observed in the control females (Figure 1, see model results in Table 1). No other main or interaction effects were significant (Table 1).

**Figure 1.**
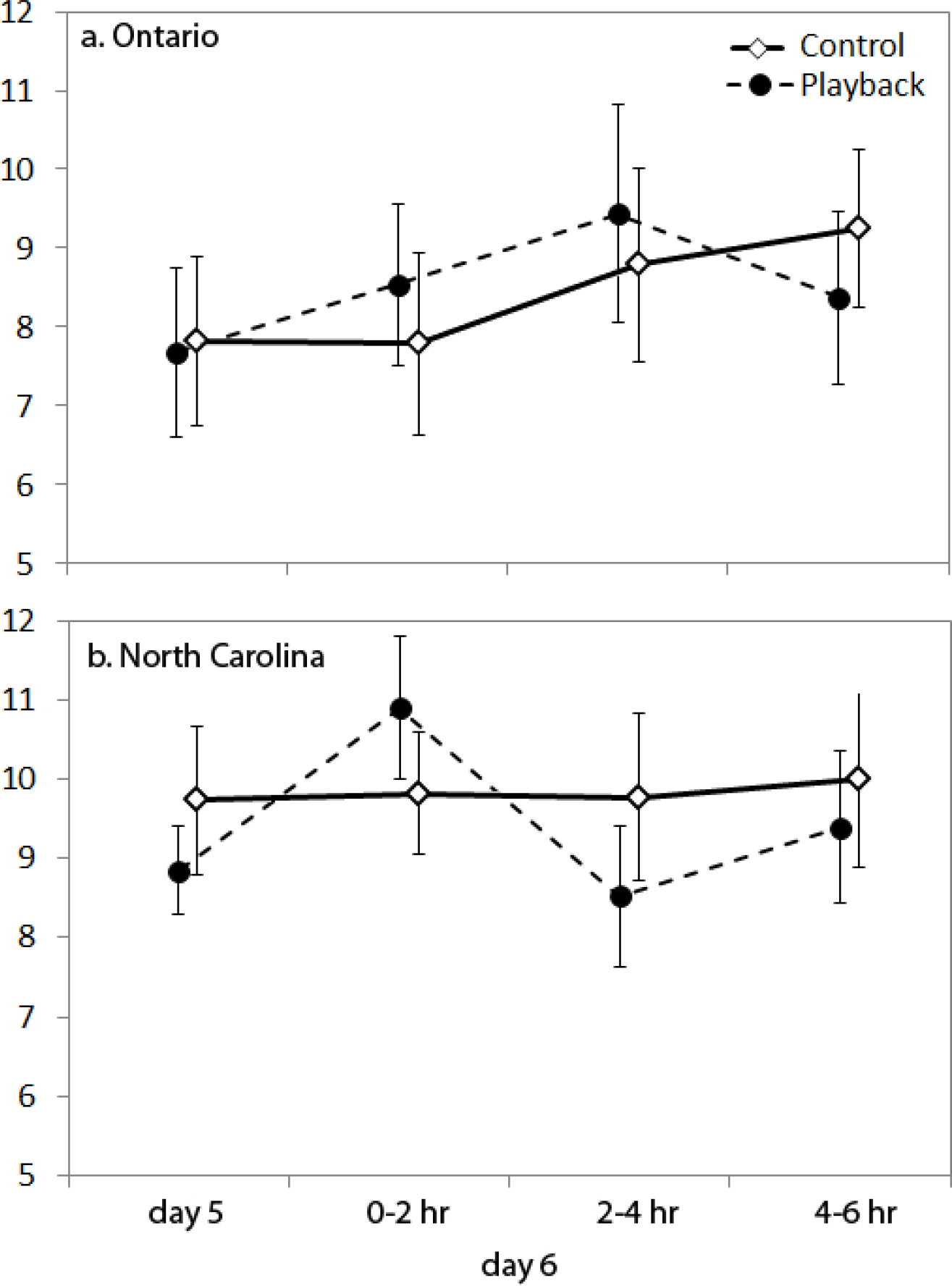
Average feeding rates of females on the day before treatment (day 5) and during the treatment (day 6), the latter in 3 two-hour increments. The feeding rates were estimated from RFID logs. The error bars denote ±1 standard error.

**Table 1.** Model averaged coefficients of predictors of female feeding rates. The comparison for the period, population, and treatment is pre-treatment (day 5), NC, and control, respectively. See text for details and supplementary materials for the specific models in the averaged model.

|  | coefficient | Estimate | Std. Error | P value | # of models |
| --- | --- | --- | --- | --- | --- |
| **Period** | during 1<sup>st</sup> 2 hrs | 0.10 | 0.06 | 0.090 | 4 |
|  | during 2<sup>nd</sup> 2 hrs | 0.03 | 0.05 | 0.568 |  |
|  | during last 2 hrs | 0.06 | 0.04 | 0.146 |  |
| **Population** | Ontario | -0.18 | 0.12 | 0.126 | 3 |
| **Period\*population** | during 1<sup>st</sup> 2 hrs*Ontario | -0.12 | 0.07 | 0.084 | 2 |
|  | during 2<sup>nd</sup> 2 hrs*Ontario | 0.11 | 0.07 | 0.102 |  |
|  | during last 2 hrs*Ontario | 0.03 | 0.06 | 0.616 |  |
| **Treatment** | Playback | -0.05 | 0.12 | 0.661 | 1 |
| **Period\*Treatment** | **during 1<sup>st</sup> 2 hrs\* playback** | **0.18** | **0.07** | **0.005** | **1** |
|  | during 2<sup>nd</sup> 2 hrs* playback | 0.03 | 0.07 | 0.625 |  |
|  | during last 2 hrs* playback | 0.01 | 0.06 | 0.886 |  |

**Table 2.**
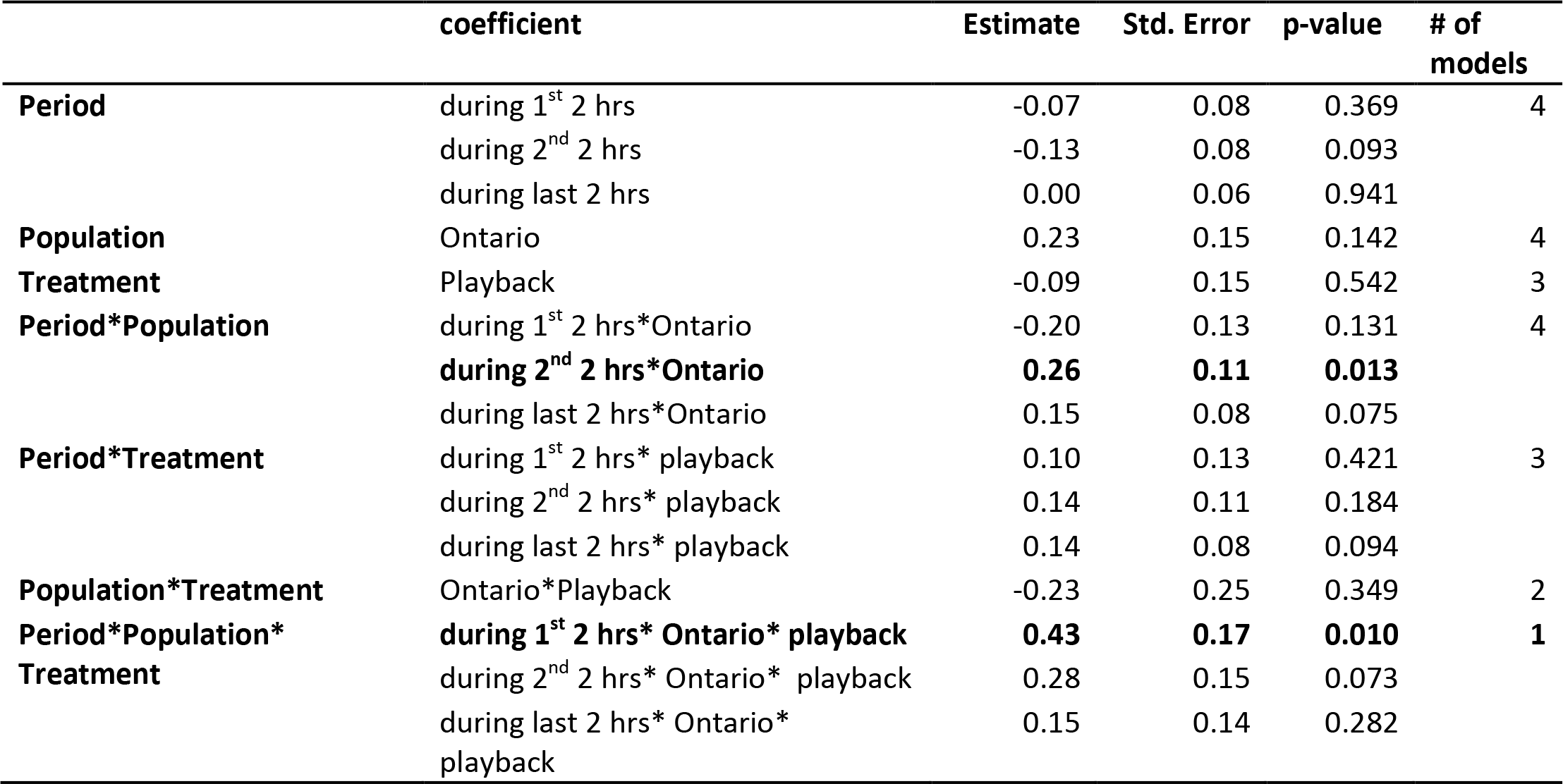
Model averaged coefficients of predictors of male feeding rates. The comparison for the period, population, and treatment is pre-treatment (day 5), NC, and control, respectively. See text for details and supplementary materials for the specific models in the averaged model.

*Corticosterone:* In the averaged model, the only significant coefficient was nesting stage: females had significantly higher cort levels during the nestling period compared to the incubation period (Figure 2). Treatment did not have a main effect or enter into an interaction with stage.

**Figure 2.**
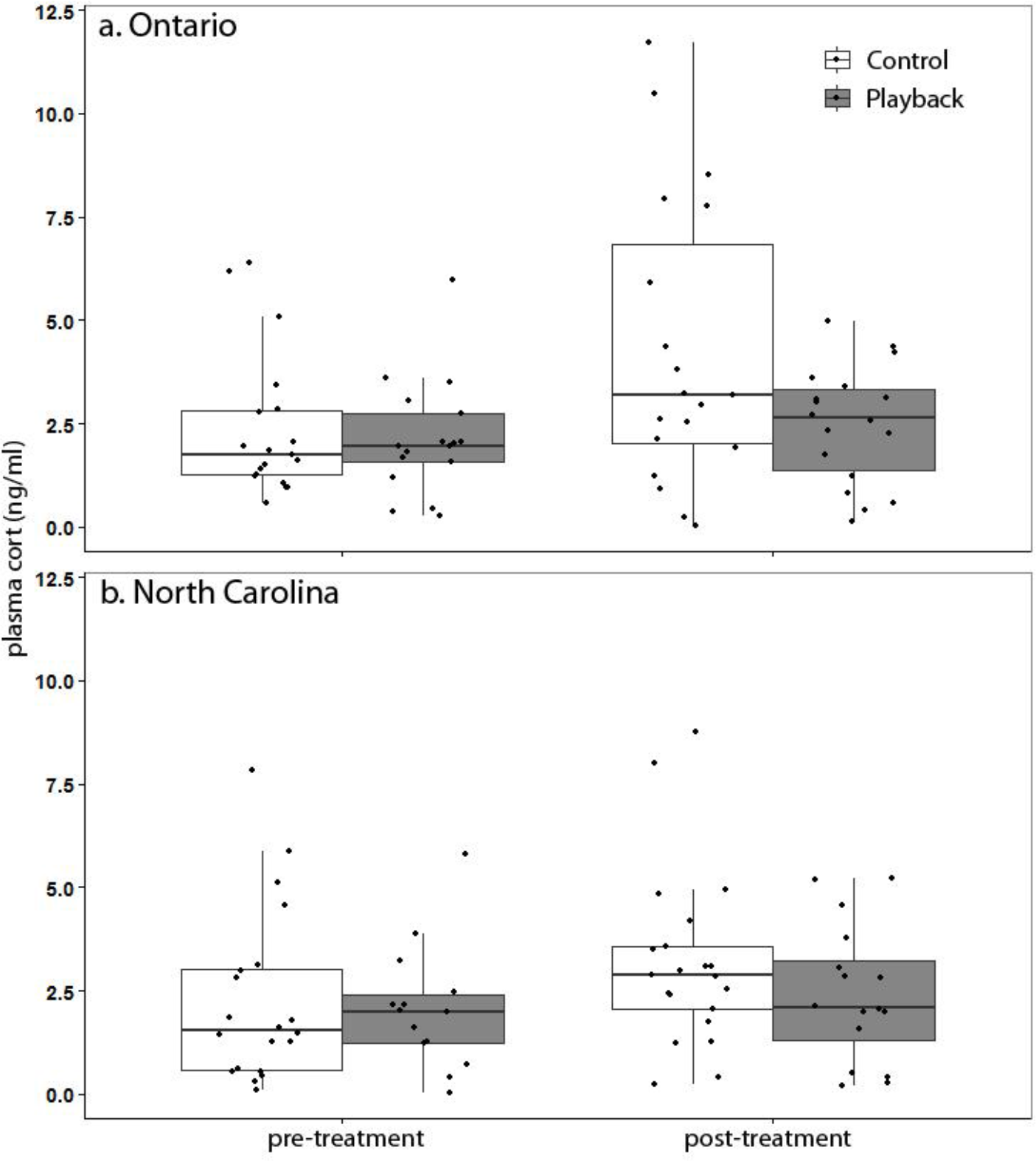
Circulating plasma cort (baseline) levels of females pre-treatment (late incubation stage) and posttreatment (immediately after the playbacks during the nestling stage). The boxplots represent the median (middle line), 25% and 75% percentiles (the lower and upper boundaries of the boxes respectively), and the 1.5 interquartile range (whiskers). Individual data points are also shown.

There was a positive correlation between female change in cort (ratio of post-to pre-treatment cort) and fledging success. In the averaged model, females with greater increases in cort from the incubation to the nestling stage fledged more offspring (Figure 3, see Table 4 for the averaged model). The effects of treatment and population were not significant.

**Figure 3.**
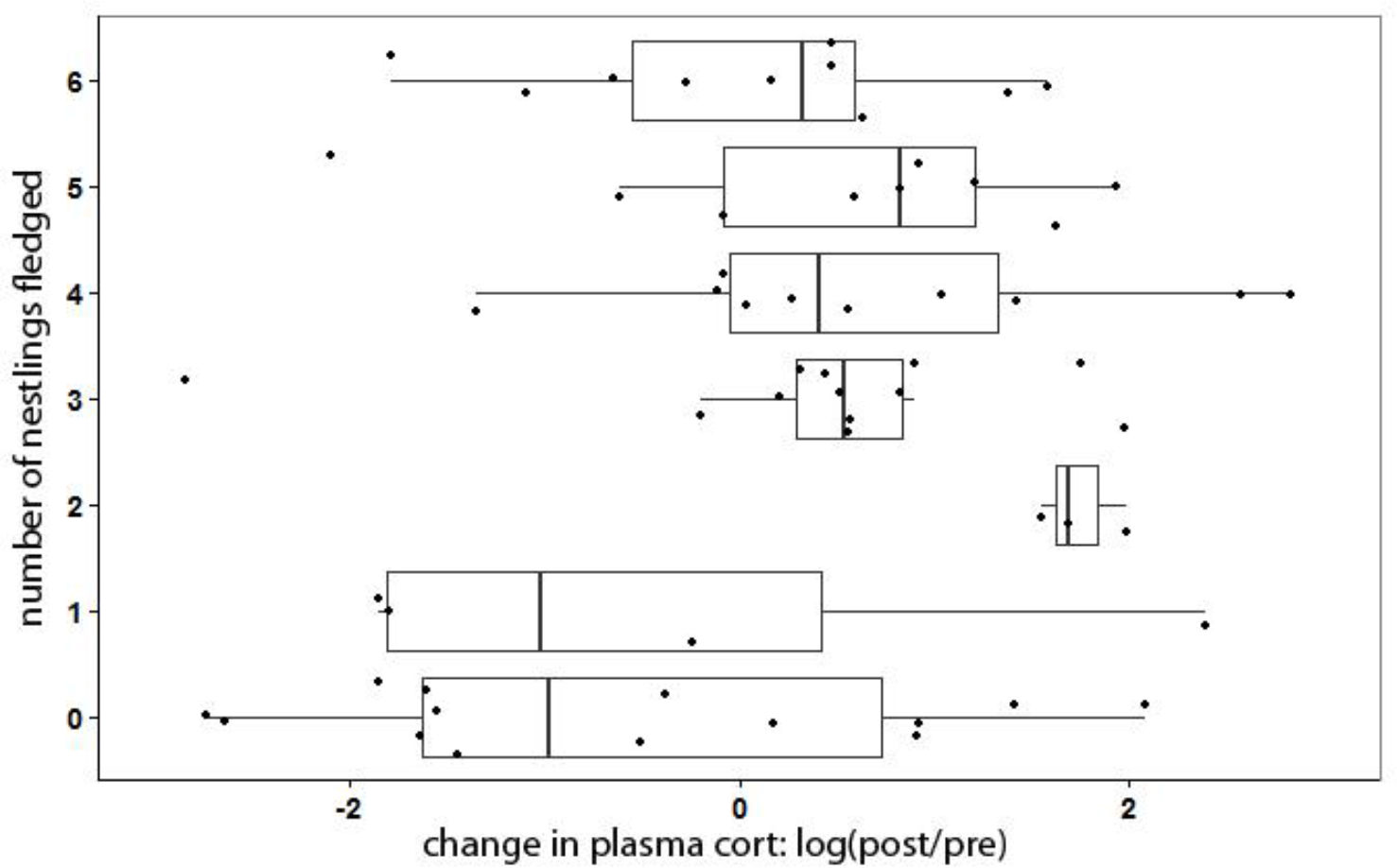
Fledging success as a function of change in baseline plasma cort levels of females from incubation to the nestling period. Change in plasma cort is depicted as the logarithm (base 10) of the ratio of plasma cort during nestling period to plasma cort during incubation. Thus, zero equals no change in plasma cort from incubation to nestling period. The boxplots show the change in cort for each number of fledging category, and the dots are individual data points.

**Table 3.**
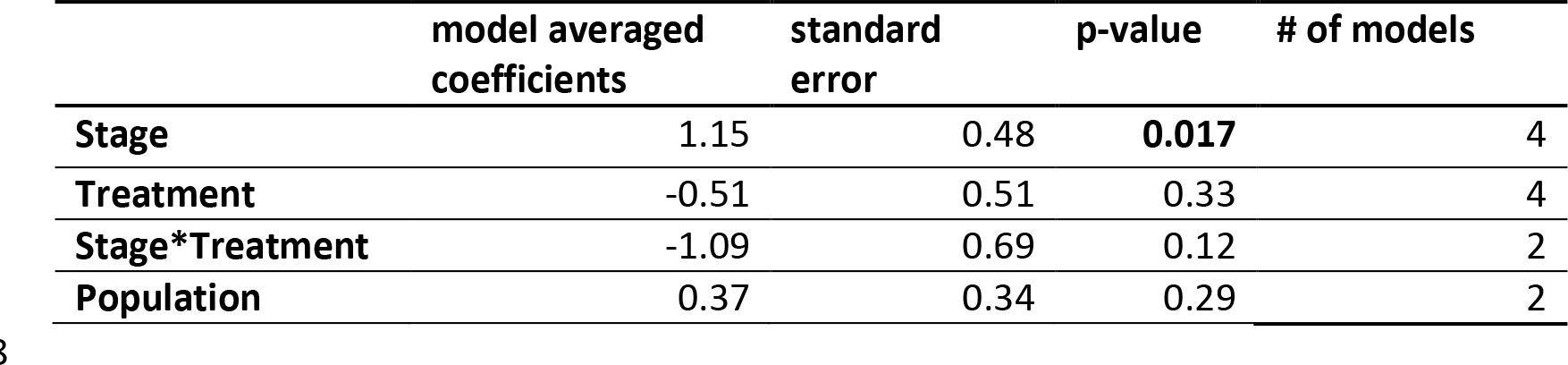
Averaged GLM model from 4 best models (within 2 ΔAlCc of best) of predictors of female CORT. The following factor levels were used as baseline for the intercept: incubation (Stage), control (Treatment), and NC (Population).Female baseline corticosterone (cort) levels significantly increased from incubation to nestling period.

|  | model averaged<br>coefficients | standard<br>error | p-value | # of models |
| --- | --- | --- | --- | --- |
| **Stage** | 1.15 | 0.48 | **0.017** | 4 |
| **Treatment** | -0.51 | 0.51 | 0.33 | 4 |
| **Stage\*Treatment** | -1.09 | 0.69 | 0.12 | 2 |
| **Population** | 0.37 | 0.34 | 0.29 | 2 |

**Table 4.** Averaged model (from 3 best models) examining predictors of the number of fledglings. For the treatment and population, control and NC were used as baseline, respectively. Change in corticosterone (cort) was positively correlated with number of nestlings fledged.

|  | model averaged<br>coefficients | standard<br>error | p-value | # of<br>models |
| --- | --- | --- | --- | --- |
| **Change in cort** | 0.10 | 0.047 | **0.029** | **3** |
| **Treatment** | 0.14 | 0.15 | 0.35 | 1 |
| **Population** | -0.08 | 0.15 | 0.61 | 1 |

*Mass:* There was a significant effect of nesting stage on body mass: almost all females lost weight from the first to second capture (see Table 5 for the best model). The main effects of population and treatment were also significant. The main effects were modified by two significant interactions: treatment by population (females in the playback treatment were lighter than the control females in the NC but not in Ontario population), and population by stage (females in NC lost more weight between the incubation and nestling periods as compared to females in Ontario). Finally, there was an effect of relative laying date with females starting to lay eggs later in the season being lighter.

**Table 5.** The best GLM model on female body mass. The model selection table revealed that the ΔAICc between the best model and the second best was 2.32. The following factor levels were used as baseline for the intercept: incubation (Stage), control (Treatment), and NC (Population).

|  | Coefficient | Std Error | p-value |
| --- | --- | --- | --- |
| Treatment | -0.83 | 0.37 | 0.03 |
| Population | -0.89 | 0.43 | 0.04 |
| Stage | -4.30 | 0.31 | <0.0001 |
| Relative Lay Date | -0.06 | 0.02 | 0.0001 |
| Treatment*Population | 1.43 | 0.53 | 0.002 |
| Population*Stage | 1.38 | 0.42 | 0.007 |

*Clutch size, nestling mass, and fledging success:* There were no significant effects of treatment on clutch size, nestling mass at day 6 and day 12, or fledging success. Nestlings in Ontario were on average significantly heavier at both day 6 and day 12 (Tables S6 and S7, Figures S2, S3), but clutch size and number of nestlings fledged did not differ between the populations. There were no significant interactions of population and treatment in any of the models (see supplementary information).

## Discussion

Our aim was to manipulate nestling begging calls to study (1) whether a perceived increase in offspring demand induces a change in parental effort of females in two different populations of tree swallows with distinct brood values. If there was a change in parental effort, we further asked whether it led to (2) an increase in baseline cort as predicted by the Cort-Adaptation Hypothesis and whether the changes in baseline cort from incubation to nestling period was predictive of fledging success. We found that (1) females increased parental effort in response to offspring begging call playback in both populations, but the increase was transient and confined to the early hours of the playback treatment. Consequently, (2) baseline cort levels obtained from blood samples at the end of the 6 hour period did not differ between the control and playback groups. However, change in baseline cort from incubation to nestling period did predict how many nestlings females fledged (with higher increases in cort associated with greater fledgling success).

### Change in parental care and baseline cort

Prior experiments on tree swallows and other species showed that glucocorticoids vary with parental effort (Bonier et al., 2009a; Bonier et al., 2009b; Bonier et al., 2011). These studies generally looked at changes in parental care over a longer period (e.g. through brood enlargement or reduction throughout the nestling period). Our aim here was to extend these findings by asking whether cort would dynamically co-vary with changes in cues of offspring demand in the short term. Our failure to detect a difference in cort between females in the playback and control groups does not support this hypothesis. Given our relatively large sample size (n = 73 nests) we do not believe the absence of a treatment effect is due to lack of power. We do however offer another caveat in that the effect of the nestling playbacks on female parental behavior was transient and confined to the first few hours of the playbacks whereas we obtained blood samples at about 6 hours after the start of the playback, and compared cort in those samples to cort measured 9-10 days prior to the playback. By the time the post-experimental blood samples were collected, female nest visit rates had returned to baseline levels, and were not significantly different between the playback and control group. Our experimental approach may have therefore lacked the precision to detect a transient increase in cort corresponding to the transient increase in female feeding rates.

The fact that playbacks had only a transient effect on female feeding behavior is somewhat surprising given that previous studies manipulating female feeding behavior through automated or manual playbacks of nestling begging calls used durations from 1 hour (in great tits, Parus major; Hinde, 2006; and in blue tits; Lucass et al., 2016) or several days (in pied flycatchers, Ficedula hypoleuca; Ottosson et al., 1997), and in both cases found an effect of the playbacks on behavior. This could reflect habituation to the stimulus, and/or an inability of the females to maintain a high rate of feeding, although the latter possibility seems less plausible given the above studies found persistent effects in different species. Whatever the cause, the transient effect of the playbacks means that any effect on cort may also have been transient and therefore would only have been detected if we had captured the females when the playbacks had their maximal behavioral effect.

The Cort-Adaption Hypothesis predicts that increases in baseline cort during the period of parental care should increase fledging success. We found support for this prediction: females with the greatest increases in cort from incubation to nestling period fledged more offspring. This finding is consistent with an earlier study in the Ontario population, in which experimentally increased broods induced greater increases in cort through the breeding season, and changes in cort within females were positively correlated with fledging success across both experimental and control groups (Bonier et al., 2011). The current results extend the previous findings by showing that the positive link between small increases in baseline cort and fledging success can be detected in natural brood sizes.

The finding that females in both populations adjusted parental care in response to begging calls is in contrast to an earlier study in this species that found effects of brood size manipulation on females consistent with the brood value hypothesis (Ardia, 2005). In that study, broods were either enlarged or reduced by 50% in two populations of tree swallows in Alaska and Tennessee. Females in Alaska increased their nest visit rate in the enlarged condition to maintain the same level of nestling condition, whereas females in Tennessee did not increase their visit rate and, subsequently, nestlings in enlarged broods were of lower condition. These results were consistent with the life-history theory and the brood-value hypothesis (Ghalambor and Martin, 2001; Martin et al., 2000). However, the present result may suggest that females in southern populations are able to increase their parental effort in the shortterm similar to their northern counterparts (the finding in the present study) but may be limited in the long term due to lower food availability in the southern populations (the situation faced by females due to permanent addition of extra nestlings in the study by Ardia, 2005). Indeed, Ardia (2005) found that insect availability was lower in the southern population, making a chronic increase in offspring demand harder to meet for the southern parents.

In summary, our results suggest that females are able to flexibly adjust their feeding rates in response to simulated increased demand from their nestlings. Additionally, increases in baseline cort levels of females from incubation to nestling period predicted fledging success across broods. Therefore, although it is unclear whether glucocorticoids are involved in short-term strategic adjustment of parental care, the data suggest that longer-term changes in baseline cort levels are positively correlated with fitness. We believe the data warrant further research into hormonal changes that may occur at shorter time scales and that glucocorticoids may play a casual role in short-term adjustment of parental effort.

## Acknowledgements

We are grateful to Alice Domalik, and Pria St John (Queen’s University), Drew Gill and Spencer Gill (Davidson College) for the excellent help in the field.

Funding: This work was supported by a U.S. National Science Foundation (NSF) grant (FB, ITM, and MFH; IOS-1145625), and by the Natural Sciences and Engineering Research Council of Canada Banting Postdoctoral Fellowship (FB). During the preparation of the manuscript, ÁZL was supported by a grant from the Hungarian Research Fund (OTKA K 113108).

